# The Abnormal Oswald Ripening of Protein Nanofiber in Myofibrillar Protein Solution

**DOI:** 10.1101/558460

**Authors:** Fuge Niu, Rui Zhang, Jiamei Fan, Weichun Pan

## Abstract

In solutions of myofibrillar protein extracted from giant squid (*Dosidicus gigas*), the size-coarsening process of protein nanofiber is complex. At high temperature (25°C), nanofiber keeps growth but with two distinguishable patterns, slow rate at the initial stage with *t^0.2^* and the fast one at the late stage with *t^2.3^*. The intersection of these two slopes is around 300 min. Meanwhile, protein concentration in solution enhances as well. These behaviors contradict to the prediction of Ostwald ripening. Thus, we call this process as abnormal. These abnormal behaviors come from the conformation change of some types of constitution protein molecules with chemical potential reduction when they dissolve from nanofiber to solution. On the other hand, low temperature (10°C) depresses this size growth. This observation suggests that temperature regulates protein molecule conformation change in nanofiber. The consequence of this abnormal Ostwald ripening process is that all protein molecules in nanofiber are redistributed. Protein molecules with the absence of conformation change in dissolution accumulate in nanofiber to cause it growing, while the rest concentrates in solution.

---

In 1896 Wilhelm Ostwald descried the phenomenon of large particle growth in the cost of small one as Ostwald ripening due to the surface tension which is proportional to the particles curvature(1-3). *As a result, the solute concentration keeps reduction throughout size-coarsening process*. Meanwhile, an essential prerequisite, no molecular conformation change when a molecule transfers from one phase to another, exists but is always ignored. This condition is satisfied in inorganic and organic compounds. However, in protein solutions containing particles with various sizes, it should be cautious to apply Ostwald ripening theory in the size-coarsening process because protein molecules are sensitive to the ambient conditions. Even salt concentration variation could induce protein molecule conformation change(4). Thus, *it is worth to verify Ostwald ripening theory in protein solutions*.

The existence of nanofiber in solutions of myofibrillar protein extracted from giant squid (*Dosidicus gigas*) is verified(5). In order to investigate its size-coarsening process in this solution, 2.73 mg mL^-1^ myofibrillar protein was diluted 10 times by the buffer solution, and was assessed by the dynamic light scattering (DLS) technique and the fluorescence spectroscopy (FS) technique immediately.

In Fig. 1a, the pattern of light intensity on time depends on the temperature. High temperature (25°C) allows the light intensity monotonically growing, while low temperature (10°C) depresses this growth with almost constant light intensity throughout the experiment. Due to high activity of enzymolysis in myofibrillar protein at 25°C(6), it is necessary to estimate its influence in this study. In Fig, 1b, the apparent concentrations of myofibrillar protein in three cases were determined via Bradford protein assay. It displaces two issues: 1) the apparent protein concentration remarkable increase overnight no matter whether ethylenediaminetetra acetic acid (EDTA) is present or absent; 2) the existence of EDTA has minor effect on the apparent concentration determinations. Since EDTA is a efficient agent to depress enzymolysis(7), it could be concluded that the effect of enzymolysis could be ignored in this study.

**Figure 1.**
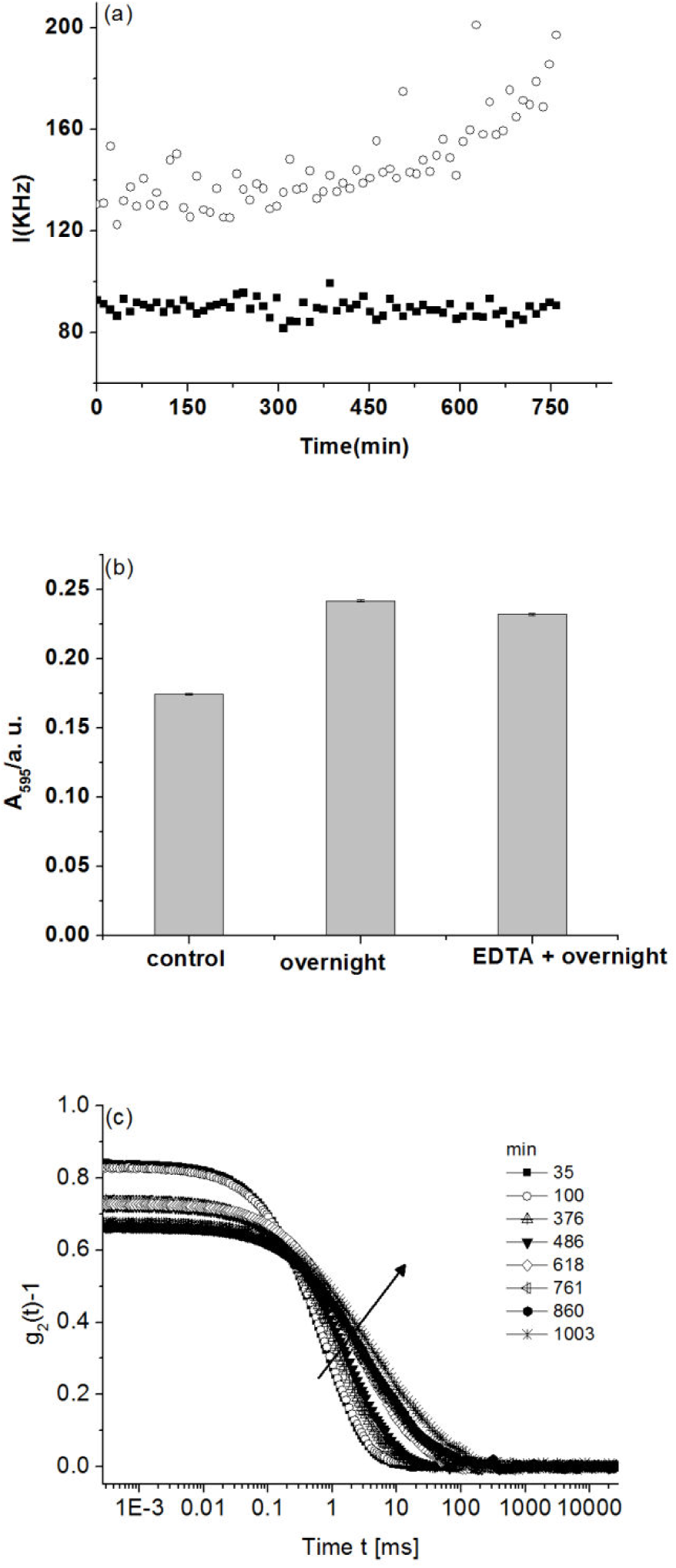
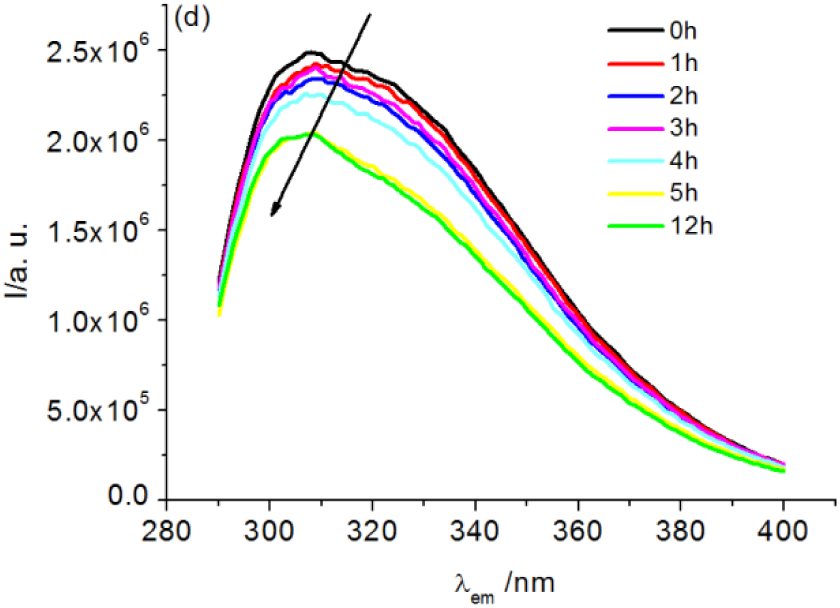
(a) Time evolutions of light intensity at various temperatures. Empty circle stands for 25°C with 0.27 mg mL^-1^ myofibrillar protein while solid square does for 10°C at the same protein concentration. (b) The influence of enzyme on protein concentration assessment in myofibrillar protein solutions. The control is 0.27 mg mL^-1^ myofibrillar protein solution. While the middle one is the solution same as control but has stayed in a heat bath at 25°C overnight; and the right one is a sample same as the middle one but added EDTA to depress enzyme activity. The Bradford method was applied for absorbance determination. (C) The autocorrelation functions of DLS at 25 °C in Fig.1a at some intervals. (d) The fluorescence emission spectrums at various intervals, which displace light intensity reduction and maximum of fluorescent spectra shifting from 309 nm to 307 nm.

Another issue of interest in Fig. 1b is the apparent concentration rising with time. Ref. 5 illustrates that the apparent concentration increase via Bradford protein assay hints the more surface of protein molecule exploring to the solvent. In other words, nanofiber dissolves into several pieces with small size. However, the last stage of light intensity observation with fast increase at 25°C in Fig. 1a is against this prediction because the light intensity is proportional to the particle number density and is six-power of the size of particle(8). The light intensity increment due to the number density increase is much smaller than its reduction due to the size shrinking during the dissolution process. In order to clarify this puzzle, the size of corresponding nanofiber was evaluated via DLS technique. Fig. 1c reveals that the decay of the autocorrelation function *g_2_(*τ*)-1* becomes slow throughout the experiment. With the aid of Laplace transform, the corresponding decay time distribution was obtained (Fig. S1). The decay time of the slow mode τ was assessed to construct Fig. S2. In contrast to the case in 10°C in which τ is almost constant throughout experiment (data do not present), the dependency of τ on the observation time has two patterns, *t^0.2^* and *t^2.3^*, with the intersection of these two slopes around 300 min (Fig. S2). In the initial stage of size-coarsening process, 0.2 is close to the value in phase-separation process, 0.212(9, 10). But in the second stage, to the best of our knowledge, 2.3 is larger than any reported value(11, 12).

Now there is a problem. On the one hand, nanofiber keeps growth. On the other hand, protein concentration in solution increases as well, which contradicts to the prediction of Ostwald ripening. How does this phenomenon occur? Hinted by the fact of tropmysin (Fig. S3) dissolution from F-actin with the expense of partial degradation of helical structure(13), we hypothesized this abnormal Ostwald ripening due to some types of protein molecule conformation change when these molecules dissolve from nanofiber to solution with chemical potential reduction. As a result, these types of protein molecules concentrate in solutions. In order to verify the hypothesis of protein molecule conformation change, FS technique was carried out immediately after 2.15 mg mL^-1^ myofibrillar protein solution was diluted 10 times. The reason to pick up FS is the high sensitivity of tryptophan to the local environment in intrinsic protein florescence. As a result, change in the emission spectrum of tryptophan is utilized to probe protein conformation change(14, 15). Especially for minor change without secondary structure modification, such as the case of this study, the traditional techniques, such as circular dichroism (CD), are infeasible. The florescent spectra of the corresponding solution at various intervals were displayed in Fig. 1d, in which the fluorescent light intensity keeps reduction with maximum of fluorescent spectra shifting from 309 nm to 307 nm. This observation indicates the occurrence of conformation change of protein molecules. In addition, the interval of remarkable fluorescent light intensity reduction, around 5 h, coincides with the intersection of the two slopes in Fig. S2.

Thus, at 25°C the slow growth rate of nanofiber at the initial stage (Fig. S1) may come from the fact of high activation energy for protein molecule conformation change, for instance, 430-490 kJ mol^-1^ for ovalbumin at pH 7^(16)^.

But what factors do dominate protein molecule conformation change? Fig. 1a hints that it is temperature rather than protein molecule concentration. When temperature is low as 10°C, nanofiber is stable, which is partially verified by the fact of animal muscles with less muscle shortening and drip loss around 10～15°C(17, 18). Only is temperature high, such as 25°C, some types of protein molecules commencing molecular conformation change when they redistribute from nanofiber to solution in myofibrillar protein. Indeed, it is found that the tropomyosin dissociation from F-actin has a minimum temperature of 35-40°C(19).

The aforementioned discussions lead to a prediction that in size-coarsening process protein molecules with the absence of conformation change when dissolving from nanofiber to solution accumulate to nanofiber while the rest types are condensed in the solution. In other words, the compositions of solution and nanofiber respectively become purer after this size-coarsening process, which was tested by the differential scanning calorimetry (DSC) assessment (Fig. 2 and Table 2), in which solutions with 2.71 mg mL^-1^ and 0.26 mg mL^-1^ respectively were assessed. The dilution operation induces the size-coarsening process in myofibrillar protein solutions. Compared with the high concentration solution, the peak is sharp in the thermogram with the low denatured temperature and less denatured enthalpy in low concentration solution. The sharp peak indicates the purer composition in nanofiber(20). In addition, the dilution operation causes nanofiber loose, corroborated by fractal dimension *d_f_* assessment with monotonic reduction (Fig S4 and Table S1). This may be the reason for low denatured temperature and less denatured enthalpy in the low concentration solution.

**Figure 2.**
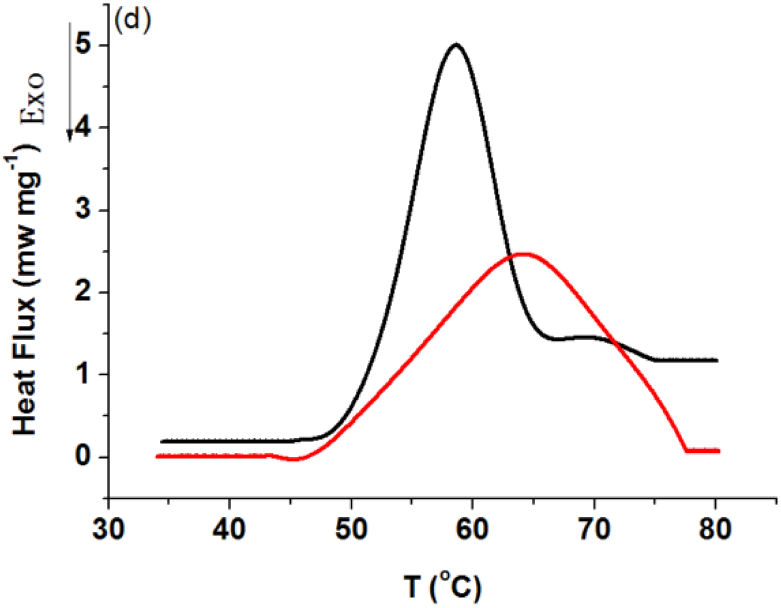
DSC determinations of myofibrillar protein solutions at two concentrations 2.71 mg mL^-1^ (black line) and 0.26 mg mL^-1^ (red line).

**Table 1.** The thermal stability of particles under various dilution ratios

| Dilution ratio | T<sub>max</sub> (°C) | ΔH (J/s) |
| --- | --- | --- |
| 1 | 63.9<sup>a</sup> | 0.529<sup>a</sup> |
| 10 | 58.6<sup>b</sup> | 0.362<sup>b</sup> |

T<sub>max</sub>, the denaturation temperature; ΔH, the endotherm enthalpy. Different letters (a-b) indicate significant (P < 0.05) difference within the same row.

From the aforementioned discussion, a crucial conclusion is made that protein molecule conformation change is an essential prerequisite for muscle protein dissolution from the solid state to solution. In addition, this conformation change process is regulated by temperature rather than protein molecule concentration. Just because many food additives have strong interactions with muscle protein molecules, they can significantly affect muscle protein molecules dissolution and thereby can influence the properties of final products made by muscle (21, 22). This observation also sheds light on clinical practices, for instance, the mechanism study of rhabdomyolysis, a complex process associated with morbidity and mortality(23).

## SUPPORTING MATERIAL

Supporting Materials and Methods, five figures, and one table are available.

## AUTHOR CONTRIBUTIONS

F. N. and R. Z. have equal contribution in this study. F. N., R. Z., and J. F. carried out the experiments. W. P. designed the study. F. N., R. Z. and W. P. analyzed the data, discussed and interpreted results. W. P. wrote the manuscript.

## ACKNOWLEDGEMENTS

The authors wish to thank Dr. Peter G. Vekilov of University of Houston for many helpful comments and criticisms. This work was supported by grants from the National Natural Science Foundation of China (31171713, 31701650), the Foundation of Food Science and Engineering, the Most Important Discipline of Zhejiang Province (JYTsp20142014), The Natural Science Foundation of Zhejiang Province (LY17C200004).

The authors declare that they have no known competing financial interests or personal relationships that could have appeared to influence the work reported in this paper.

## SUPPORTING CITATIONS

References (24–30) appear in the Supporting Material.

