## supporting materials for "The Abnormal Oswald Ripening of Protein Nanofiber in Myofibrillar Protein Solution"

**Raw materials**

Dead giant squid stored at −20°C for less than 3 months after being caught and immediately being frozen without any further treatment, were obtained from the Zhoushan Second Marine Fisheries Company in Zhejiang, China. All squid (usually four organisms of average mantle size 1 m) were deheaded, degutted, and cleaned using water after they were defrosted in air to room temperature. The mantles with the fins were bagged and stored at–80 °C for further experiments.

$CH_3COOH$, $H_3PO_4$, $Na_2HPO_4$, $NaH_2PO_4$, KCl, KI, NaOH, HCL, sodium potassium tartrate, Ethylenediaminetetra acetic acid (EDTA) and $CuSO_4$•$5H_2O$ with analytical grade were provided by Jincheng (Hangzhou, China); Triton-X100 (Chemical pure) was obtained from Aladdin (Shanghai, China); Coomassie brilliant blue (G-250) and bovine serum albumin (BSA) were purchased from Amresco (Cuyahoga, OH, USA).

All water was obtained from Millipore-Q (Merck, Burlington, MA, USA).

**Preparation of myofibrillar protein**

Myofibrillar protein samples were prepared following the procedure described by Hashimoto, Watabe, Kono, and Siro(1) with minor modification.

The ground muscle mixture with buffer A (0.1M KCl，0.02 M Tris-HCl，pH7.5) (1:6, w/v) was homogenized using a tissue homogenizer (Homogenizer T25 basic, IKA, Staufen, Germany) at a speed of 9600 *rpm*. After this, the connective tissues were removed by filtration using two layers of gauze. The filtrate was centrifuged at 5000 *g* for 15 min at 4°C (Biofuge Stratos, Thermo Scientific Inc., Belmont, CA, USA). The supernatant was removed and the sediment was suspended with 2 volumes of phosphate buffer B (0.1 M KCl，0.02 M Tris-HCl，1%Triton(w/v)). This sample was again centrifuged at 5000 *g* for 15 min at 4°C (Biofuge Stratos, Thermo Scientific Inc., Belmont, CA, USA). The pellet (the myofibrillar protein precipitate) was washed with 8 volumes of buffer A. The mixtures were centrifuged at 5000*g* for 15 min at 4°C. The supernatant was removed and the washing procedure was repeated thrice. The final pellets were placed in tubes and stored in a refrigerator at 4°C. All pellets were used within 24 h.

An aliquot of 10g of myofibrillar protein precipitation was mixed with 20mL of 0.02 M Tris-HCl buffer solution (0.5 M KCl, pH 6.5) at 4°C by stirring for 3 h then centrifuged at 5000 *g* for 10 min at 4°C. The supernatant was collected as a sample solution with initial concentration (ICS). The solutions with various protein concentrations were achieved by the dilution of aliquot of ICS with 0.02 M Tris-HCl buffer solution (0.5 M KCl, pH 6.5) by integer times.

In order to assess the stability of floating protein nano-fibre in solutions, the protein solutions with various dilution ratios were prepared and were covered by parafilm to prevent dust dropping in and water evaporation. The solutions were put in a heat bath (DF-101S, Yuhuayiqi, Gongyi, China) at 25°C and 10°C respectively overnight.

It is well known that muscle of giant squid contains enzymes high biological activity with the result of myosin molecule degradation and other protein molecule autolysis(2), thus it is worth to probe the effect of these enzymes on protein concentration determination via EDTA because of its inhibition on these enzymes' activity in giant squid(2). The same solutions mixed with EDTA (1 mM) were treated with the same procedure.

**Myofibrillar protein concentration assessment**

Since almost all available methods for protein concentration determination have shortcoming when they apply in myofibrillar protein solutions, we picked up Bradford assay(3). 1mL protein solution was mixed with 5mL Coomassie Brilliant Blue reagent (0.01% (w/v) Coomassie Brilliant Blue G-250, 4.7% (w/v) CH3COOH, and 8.5% (w/v) H3PO4). Waiting for 5 min, the mixture was assessed at 595 nm via a spectrophotometer (UV-2600, Shimadzu, Kyoto, Japan) in triplicate at room temperature. Figure S5 is the standard curve of bovine serum albumin.

**Light scattering measurement**

Dynamic light scattering experiments were performed using an ALV/CGS-3 goniometric system (ALV, Langen, Germany) with a He–Ne laser ($\lambda$ = 632.8 nm) as well. The detector angle was 90$^{o}$ and the acquirement time for each shot was 1 min. The measurement temperature was 25$^{o}$C and 10$^{o}$C respectively. A cuvette containing ICS was inserted to a holder of ALV system after the temperature stability was achieved. And in every 11 min, the intensity of light scattering of ICS had being determined with acquirement time of 1 min in triplicate for 24 h.

The intensity correlation function $g_2(\tau)$ was expressed(4):

$$g_2(t) - 1 = (D_1 e^{-t/\tau_1} + D_2 e^{-t/\tau_2})^2 + \varepsilon \qquad (2)$$

where: D is the amplitude of light intensity; $\tau$ is the characteristic time of the diffusion of the scatterer, determined by numerically inverting the Laplace transform of $(g_2(\tau) - 1)^{1/2}$ with the aid of the software package provided by ALV-Gmbh based on CONTIN algorithm(5, 6); and $\varepsilon$ is noise.

**Fluorescence spectra**

All fluorescence spectra measurements were performed using a fluorescence spectrometer (FLS980, Edinburgh Instrument, France). The excitation wavelength was set at 280.00 nm, and the emission was monitored within the range of 290.00-400.00 nm, using a fixed exit and emits slit width of 2.00 nm. The scan step is 1.00 nm with the dwelling time 0.20 sec. All analyses were carried out in triplicate using separate samples.

**Differential scanning calorimetry (DSC) assessment**

ICS and the solution of ICS with 10 time dilution were assessed by a calorimeter (C80, Setaram, Caluire, France). The protein solution with a mass of ~4.000 g was put into a sample cell, while the same amount of a 0.02 M Tris-HCl buffer solution was added into a reference cell. Before and after each assessment, the masses of each cell were weighted by a balance (ME204E, Mettler Toledo, Columbus, OH, USA) in order to prevent the error due to water evaporation during the measurement with the tolerance of mass loss < 0.02%. The temperature range spread from 35°C to 100°C with the heating rate of 1°C min-1. The final results were treated via the software Calisto offered by Setaram.

**The fractal dimension $d_f$ assessment**

The measurement angle spread from 30° to 150° with 2° a step. Three shots were carried out at each angle with the acquirement time of 1 min at 25°C. Based on the expression $I \propto q^{-d_f}$, $d_f$ is determined by the linear fitting when the natural logarithm scales were utilized for both variables(7).

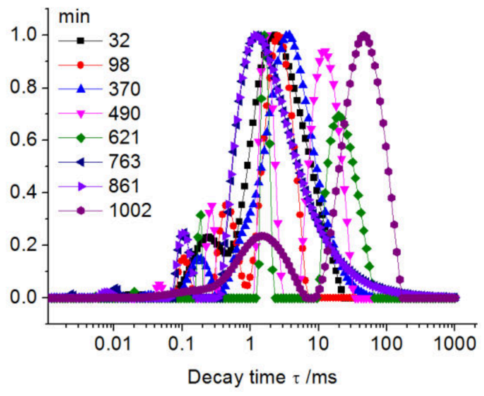


Figure S1. The corresponding decay time distributions of Fig. 1a

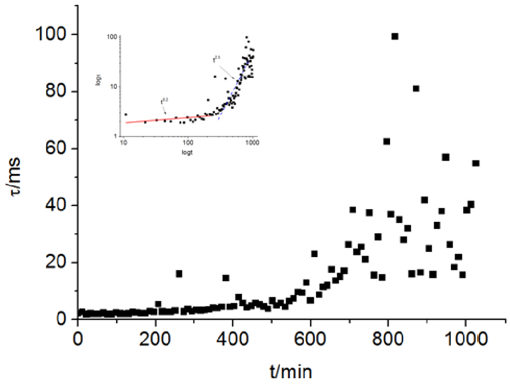


Figure S2 The corresponding decay time evolution of Fig. 2a. The inset figure is a re-plot with log-log scale, which displaces the pattern of two-part differentiated by the protein nano-fibre growth rate. The first part is characterized by the slow growth with slope of 0.2 while the second part by fast with the slope of 2.3. The red and blue lines are guided for eye.

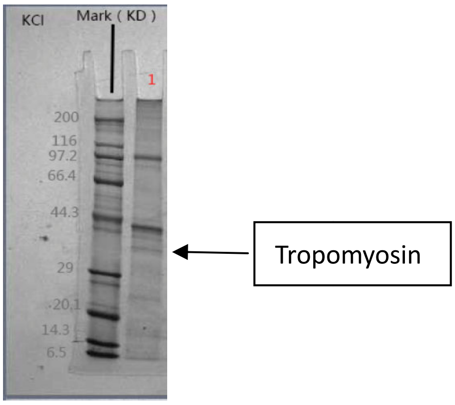


Figure S3 The SDS-PAGE analysis. The different lanes present the myofibrillar protein solution in various KCl concentrations.

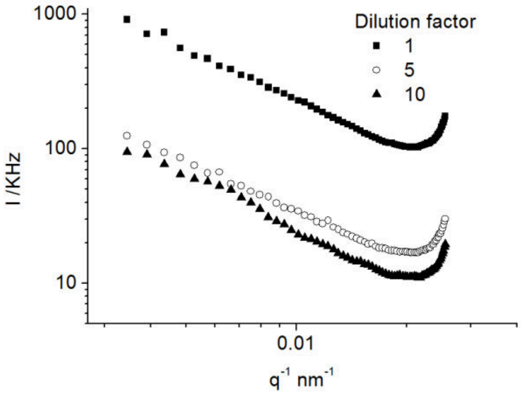


Figure S4 The scattering light intensity dependency of q at various dilution factors.

Table S1 The fractal dimensions of nanofiber at various dilution factors.

| Dilution ratio | $d_f$ |
|---|---|
| 1 | 1.265[a] |
| 5 | 1.208[b] |
| 10 | 1.087[c] |

Different letters (a-c) indicate significant (P < 0.05) difference within the same row.

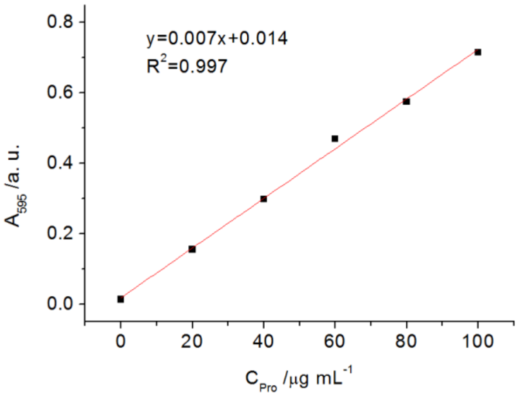


Figure S5. The standard curve of bovine serum albumin.
